# CRISPRpred(SEQ): a sequence based tool for sgRNA on target activity prediction [(almost) beating Deep Learning pipelines by traditional machine learning]

**DOI:** 10.1101/655779

**Authors:** Ali Haisam Muhammad Rafid, Md. Toufikuzzaman, Mohammad Saifur Rahman, M. Sohel Rahman

## Abstract

An accurate and fast genome editing tool can be used to treat genetic diseases, modify crops genetically etc. However, a tool that has low accuracy can be risky to use, as incorrect genome editing may have severe consequences. Although many tools have been developed in the past, there are still room for further improvement. In this paper, we present CRISPRpred(SEQ), a sequence based tool for sgRNA on target activity prediction that leverages only traditional machine learning techniques. We compare the results of CRISPRpred(SEQ) with that of DeepCRISPR, the current state-of-the-art, which uses a deep learning pipeline. In spite of using only traditional machine learning methods, we are able to beat DeepCRISPR for the three out of four cell lines in the benchmark dataset convincingly (2.174%, 6.905% and 8.119% improvement for the three cell lines), which is quite outstanding.

## 1 Introduction

Genome-editing technology has become extremely popular in recent times and more and more works are being done on it everyday. One of the more widely used genome-editing technologies is CRISPR-Cas9 (Clustered Regularly Inter-spaced Short Palindromic Repeats-CRISPR-associated protein 9). CRISPR-Cas9 is preferred over other technologies because of its higher degree of flexibility and accuracy in cutting and pasting genes. It is also more cost efficient than other methods. In addition, it allows to remove more than one gene at a time. By using CRISPR-Cas9 we are now able to manipulate multiple genes in plant and animal cells within weeks which would otherwise have taken years before. Moreover CRISPR-Cas9 can also edit genes of those species which were once considered resistant to genetic engineering.

### 1.1 Motivation

CRISPR-Cas9 is already being applied in different fields ranging from agriculture to human health. In agriculture, we can design new disease resistant [1] higher yielding crops. By editing germ-line cells, the medical issues such as infertility and other genetic diseases can be treated. One of the fascinating prospects is the creation of transgenic animals to harvest human organs. There is also possibilities of gene therapy that aims to insert normal genes into the cells of people who suffer from genetic disorders such as cystic fibrosis, hemophilia or Tay-Sachs diseases. But due to the off target effect, use of CRISPR-Cas9 on human is still considered a risk [2].

### 1.2 Previous Works

In gene editing with CRISPR we use a single guide RNA(sgRNA) with Cas-9 protein. The cut position in the DNA is specified by that sgRNA. Theoretically we can engineer the sgRNA so that it binds to the site where it exactly matches the complement of the DNA strand. But in practice, cutting efficacy may vary significantly [3, 4, 5]. For this reason predictive models are essential in designing sgRNA.

There are many tools available for designing sgRNAs. These tools differ in the type of models used, selected features, genomes etc. The tool sgRNA Designer [6] followed the rules proposed by root laboratory [7]. Their training dataset contained genes from human and mouse cells. They used Support Vector Machine (SVM) [8] classifier to select the best subset of features from among the 586 available features. Finally a logistic regression model was trained for prediction [9]. Later this dataset was enriched and used by CRISPRpred [10], E-CRISPR [11], PROTOSPACER [12], CHOPCHOP [13] and WU-CRISPR [14]. Among these tools, CRISPRpred performed significantly well than others. The authors in [10], incorporated position specific and position independent features ranked by random forest [15, 16] and trained a SVM model for prediction.

DeepCRISPR is the current state-of-the-art tool in this domain which has been proposed very recently in [17]. DeepCRISPR has first leveraged a deep unsupervised representation learning strategy to train a Deep Convolutionary Denosing Neural Network (DCDNN) based Autoencoder [18] for learning features. These features are then fed into a Convolutional Neural Network (CNN) for training the prediction model. To evaluate DeepCRISPR, the authors in [17] have used a dataset comprising sgRNAs from four different cell types: HCT116, HEK293, HeLa and HL60. DeepCRISPR achieved an ROC-AUC score of 0.874, 0.961, 0.782 and 0.739 for the four cells respectively, beating the previous tools.

### 1.3 Our Contributions

In this paper, we present CRISPRpred(SEQ), a tool to predict on-target activities of sgRNAs using a traditional machine learning pipeline. A characteristic feature of CRISPRpred(SEQ) is that, unlike the previous models, it only focuses on sequence based features. This is motivated by the empirical assertion of the natural belief (please see the recent Ph.D. thesis of Rahman [19] and the published results thereof in [20, 21, 22]) that the functional and structural information of a biological sequence are intrinsically encoded within its primary sequence. Indeed, our results in this research work further strengthen this assertion empirically as CRISPRpred(SEQ) has performed exceptionally well and has almost beaten the state-of-the-art deep neural networking pipeline (i.e., DeepCRISPR) leveraging only traditional machine learning techniques and focusing only on primary sequence based features. In particular, CRISPRpred(SEQ) has improved upon the results of DeepCRISPR by 2.174%, 6.905% and 8.119% for the cells HCT116, HeLa and HL60 respectively. This seems remarkable for at least two important reasons as follows. Firstly, this suggests that traditional machine learning techniques may have the potential to compete at par with the deep learning techniques, which is an interesting result in itself. And secondly, this further suggests that our (human) feature engineering exercise, as has been done in case of CRISPRpred(SEQ), which is an integral step in any traditional machine learning pipeline, has been able to beat automatic feature extraction of the deep networking pipeline (in this case, a Convolutional Neural Network (CNN)) of DeepCRISPR which was further enhanced by a pre-feature extraction step using a deep unsupervised learning leveraging a DCDNN based Autoencoder. Finally, unlike the experimental setup of DeepCRISPR, our experimental setup is unfortunately limited by computational resources and CRISPRpred(SEQ) promises to do even better if more computational resources can be leveraged. We plan to make CRISPRpred(SEQ) publicly available as an web-based tool. For now, we will provide the tool on request.

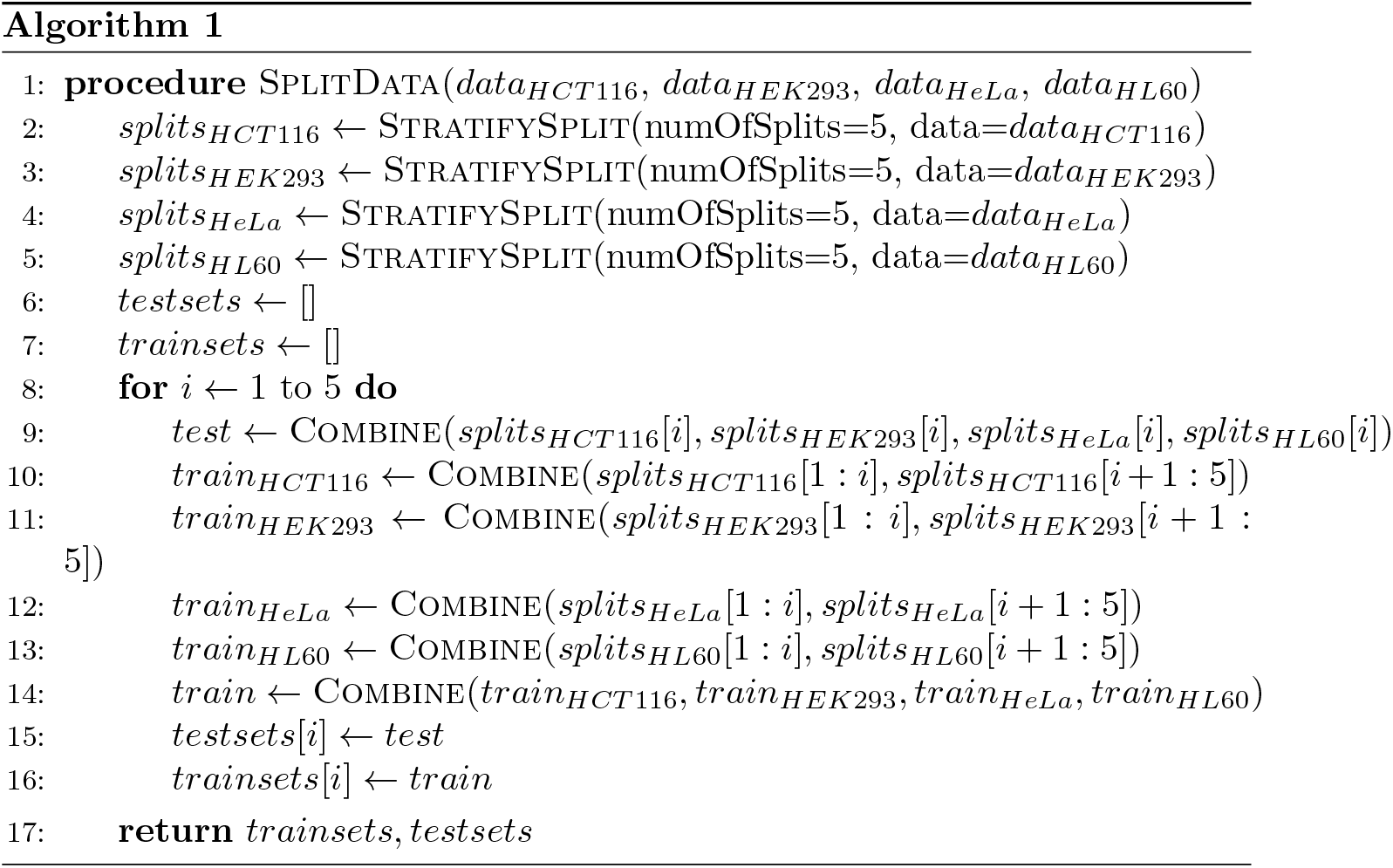

## 2 Results

### 2.1 Datasets

For all our experiments we used the on-target dataset used in DeepCRISPR which was also used by Haeussler et al. [23]. This whole dataset contains a total of 16749 labeled sgRNAs for four cell types, namely, HCT116, HEK293, HeLa and HL60. We carried out our experiments in 3 different settings (A, B and C) to be described below.

We have split the dataset for four cells separately in 5 parts where the splits were stratified by data labels. In turn, we used the combination of one part of each cell as the testing dataset (20% of the data) and the combination of other 4 parts of each cell were used as the training dataset (80% of the data). So, each part was used as the testing data while the remaining parts were used as the training data. This procedure is shown in Algorithm 1 for better understanding.

### 2.2 Results of Experimental Setup A

In this setting, we have used the training data to train the pipeline used in CRISPRpred [10] (CRISPRpred(SEQ)-A). We have extracted position independent and position specific features. After that, we have used random forest to rank the features using gini score [24] and then selected the top 2899 features. Then support vector machine has been used to train the final model.

The results were compared with DeepCRISPR [17], sgRNA Designer [6], SSC [25], CHOP-CHOP [13], CRISPR MultiTargeter [26], E-CRISP [11], sgRNA Scorer [27], Cas-Designer [28] and WU-CRISPR [14] (Figure 1). Following the paper presenting DeppCRISPR [17], we have used ROC-AUC as the metric for comparison.

**Figure 1:**
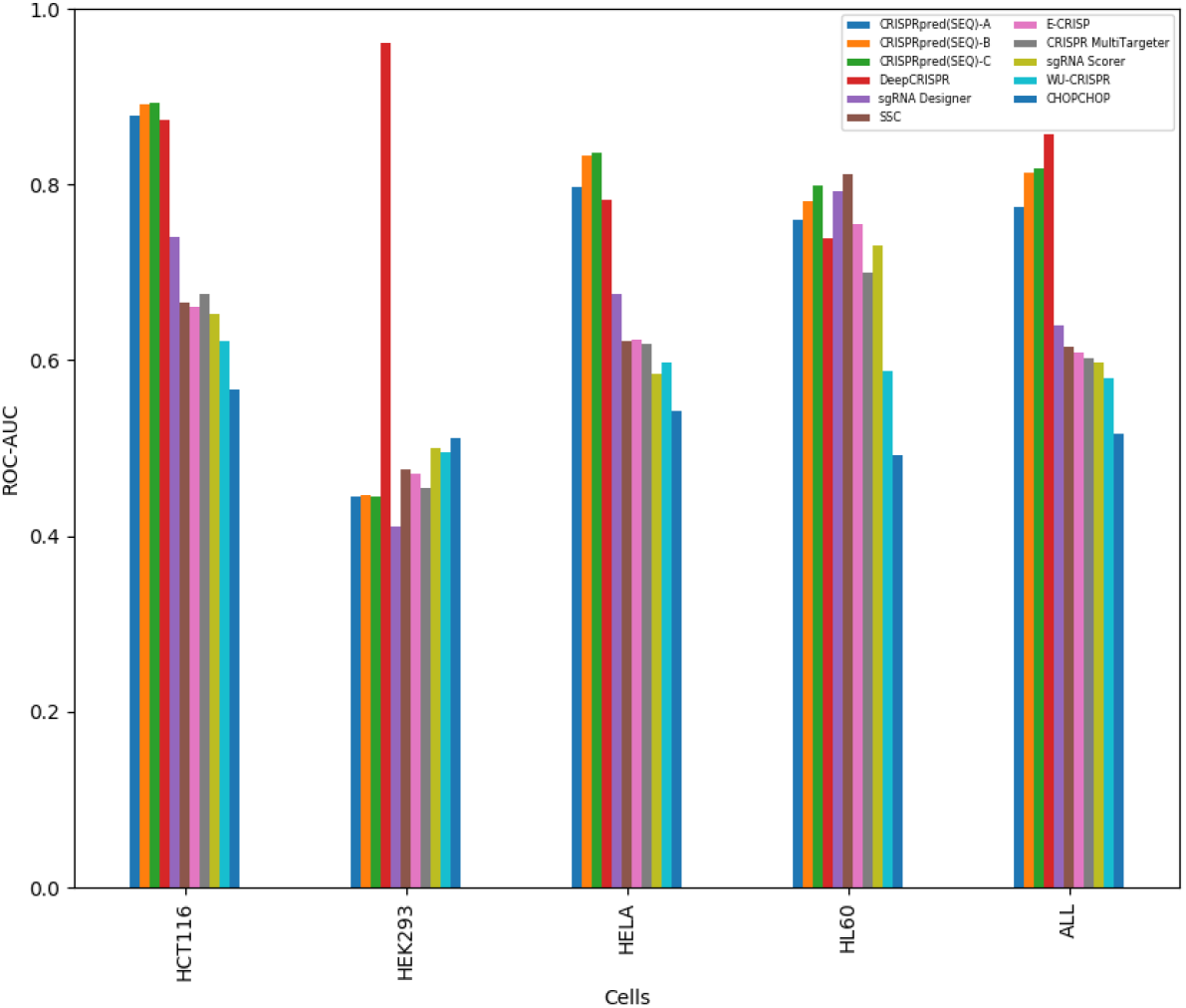
Comparison of performance of various tools in all three experimental settings (A, B, C). Y-axis denotes the ROC-AUC and X-axis denotes the cell types. In all three settings, CRISPRpred(SEQ) has convincingly beaten DeepCRISPR in 3 out of 4 cells, i.e., in HCT116, HeLa and HL60. However, in HEK293, DeepCRISPR performs far better than CRISPRpred(SEQ) (please also see a relevant discussion in the Discussion section). CRISPRpred(SEQ)-C performs slightly better than CRISPRpred(SEQ)-B which in turn outperforms CRISPRpred(SEQ) in all cell lines.

From the results we observe that the state of the art tool, DeepCRISPR, has achieved an ROC-AUC score of 0.874, 0.961, 0.782, 0.739 for the cells HCT116, HEK293, HeLa and HL60 respectively. CRISPRpred(SEQ)-A has achieved an ROC-AUC score of 0.879, 0.444, 0.797 and 0.759 for the four cells respectively. Thus, CRISPRpred(SEQ)-A was able to beat DeepCRISPR for 3 out of 4 cells.

### 2.3 Results of Experimental Setup B

In this setting, unlike the normal CRISPRpred pipeline, we have used extremely randomized trees instead of random forest to rank and select the features. Like before, the features are extracted and then ranked using gini score. Then we have selected the features having a score greater than or equal to the mean gini score, thereby selecting around 1995 features. Then we have performed 3 fold cross validation on the training data for tuning the SVM parameters *C* and *γ*. Increasing the value of *γ* means that SVM tries more to exactly fit the training data. On the other hand, the *C* parameter controls the smoothness of the decision boundary. We have tuned our model for *C* values of 1, 10 and 100 and *γ* values of 0.0001, 0.001, 0.01. We have achieved the best cross validation result for *C* =10 and *γ* = 0.001. We have determined the best hyperparameters based on the metric ROC-AUC. The detailed results of cross validation is given in Table 1.

**Table 1:** The results of 3 fold cross validation hyperparameter tuning of Experiment B. All the values in the table are ROC-AUC. The best result has been achieved for *C* =10 and *γ* = 0.001

| $C \backslash \gamma$ | 0.0001 | 0.001 | 0.01 |
| --- | --- | --- | --- |
| 1 | 0.702 | 0.775 | 0.759 |
| 10 | 0.733 | 0.781 | 0.758 |
| 100 | 0.765 | 0.781 | 0.758 |

Subsequently, We have retrained the model (CRISPRpred(SEQ)-B) using the best hyperparameters and then have compared the results with the previous tools as is done in Experiment A (Figure 1). CRISPRpred(SEQ)-B has achieved an ROC-AUC of 0.892, 0.446, 0.832 and 0.781 for the cells HCT116, HEK293, HeLa and HL60 respectively, improving further upon the results of CRISPRpred(SEQ)-A.

### 2.4 Results of Experimental Setup C

In this setup, we have added a new feature type called n-gapped dipeptide along with the previous features used in Experiment A and B. Similar to the previous pipeline, then the features have been ranked and selected using extremely randomized tree, thereby selecting a total of around 1957 features. We again have performed 3 fold cross validation to tune the SVM parameters *C* and *γ*. The results of hyperparameter tuning are presented in Table 2. After the final training of the model (CRISPRpred(SEQ)-C) with the best hyperparameters (C = 10 and *γ* = 0.001), we have compared the results with the previous tools (Figure 1). CRISPRpred(SEQ)-C registers a slight improvement over CRISPRpred(SEQ)-B with an ROC-AUC of 0.893, 0.445, 0.836 and 0.799 for the cells HCT116, HEK293, HeLa and HL60 respectively, which can be attributed to the newly added n-gapped dipeptide features.

**Table 2:** The result of 3 fold cross validation hyperparameter tuning of Experiment C. All the values in the table are ROC-AUC. The best result is achieved for *C* =10 and *γ* = 0.001

| $C \backslash \gamma$ | 0.0001 | 0.001 | 0.01 |
| --- | --- | --- | --- |
| 1 | 0.705 | 0.783 | 0.763 |
| 10 | 0.732 | 0.788 | 0.762 |
| 100 | 0.763 | 0.788 | 0.762 |

## 3 Discussion

CRISPRpred(SEQ) has performed remarkably well for 3 out of 4 cells for the DeepCRISPR dataset thereby (almost) beating a deep learning pipeline (DeepCRISPR) leveraging only classical machine learning methods. However, the performance for HEK293 cell is not up to the mark; thus a brief discussion on this point is in order.

HCT116 is a human colon cancer cell line used in therapeutic research and drug screenings [29]. HeLa cell line was derived from cervical cancer cells [30] and HL60 cell line is a human leukemia cell line [31]. On the contrary, HEK293 are a specific cell line originally derived from human embryonic kidney cells grown in tissue culture [32]. Thus, on the surface, it seems that, the features we have designed are capable of capturing the characteristics of cancer cells but do not work well for HEK293 cell, which is not a cancer cell.

At this point, we also would like to make a remark on the (excellent) result of DeepCRISPR for HEK293 cell line. As has been mentioned above, DeepCRISPR has first leveraged a deep unsupervised representation learning strategy to train a DCDNN based Autoencoder [18] for learning features. Here they have used over 70GB of unlabeled data by generating all 23 nucleotide sequences ending with NGG, NAG, CCN and CTN from Human genome directly and then using ENCODE data to add epigenomic information to each record [33]. This resulted in about 0.68 billion unlabeled sgRNA sequences. Now, since HEK293 is a specific cell line originally derived from human embryonic kidney cells, this raises the question whether the features generated above from human genome directly has given some extra biases/advantages to the pipeline particularly for HEK293. Extra bias maybe introduced if the overlapping sgRNA between test data and unsupervised data was not removed.

Finally we note that, apart from some important changes in the machine learning pipeline employed, CRISPRpred(SEQ) principally differs from its predecessor, CRISPRpred, in that the former only focuses on sequence based features whereas the latter have considered other types of features as well.

## 4 Methods

First, we preprocessed and examined the dataset we were going to use. After that, we extracted the designed features from the dataset. Then we ranked and selected features from it by using extremely randomized trees. Finally, we tuned the hyperparameters of SVM and finally trained the prediction model. The steps are described below in details and the method pipeline is shown in figure 2

**Figure 2:**
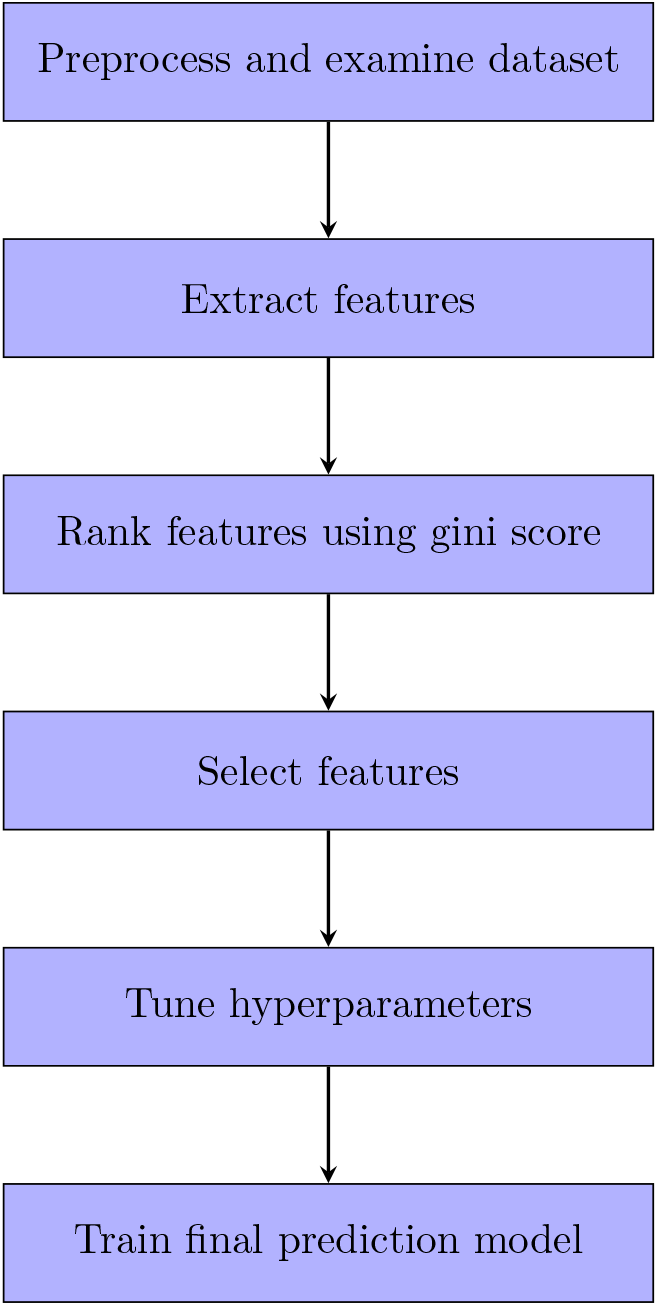
Training pipeline, the steps of building the final prediction model.

### 4.1 Dataset

The dataset used consists of sgRNAs from four cell types: HCT116, HEK293, HeLa, HL60. This dataset has been used in [17] and [23] before. There are 4239 sgRNAs from HCT116, 2333 sgRNAs from HEK293, 8101 sgRNAs from HeLa and 2076 sgRNAs from HL60, amounting a total of 16749 sgRNAs with experimentally validated known knockout efficacies from 1071 genes [17] genes.

We examined how the normalized counts of the nucleotides vary from cell to cell for this dataset (Figure 3). The position wise distribution of nucleotides in sgRNAs for four cells are also shown in Figure 4.

**Figure 3:**
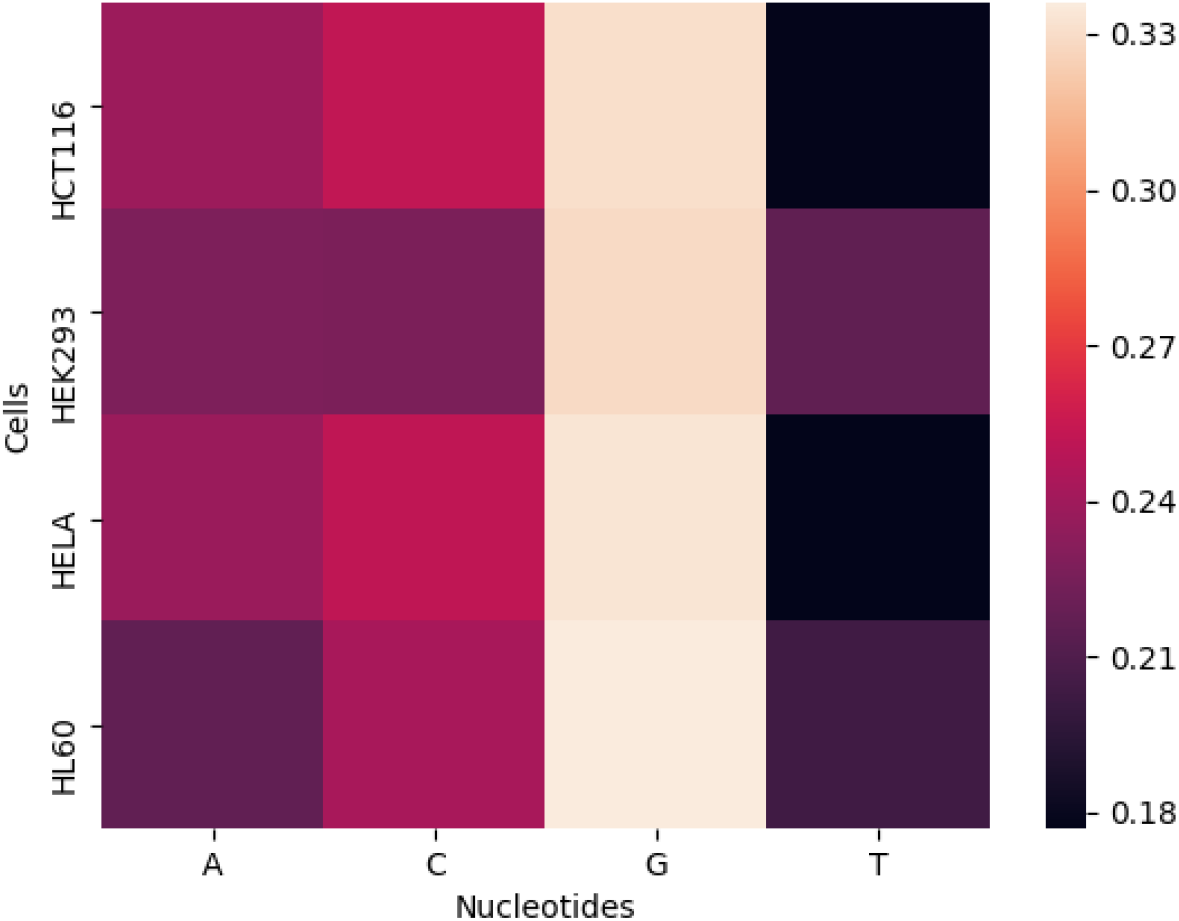
Cell wise nucleotide distribution in sgRNAs which shows the normalized count of nucleotides Adenine (A), Cytosine (C), Thymine (T) and Guanine (G) for 4 cell types: HCT116, HEK293, HeLa and HL60. Adenine and Cytosine appears more in HCT116 and HeLa cell sgRNAs. Thymine count is greater in HL60 and HEK293 cell types. The count distribution of Guanine does not change considerably across the cells

**Figure 4:**
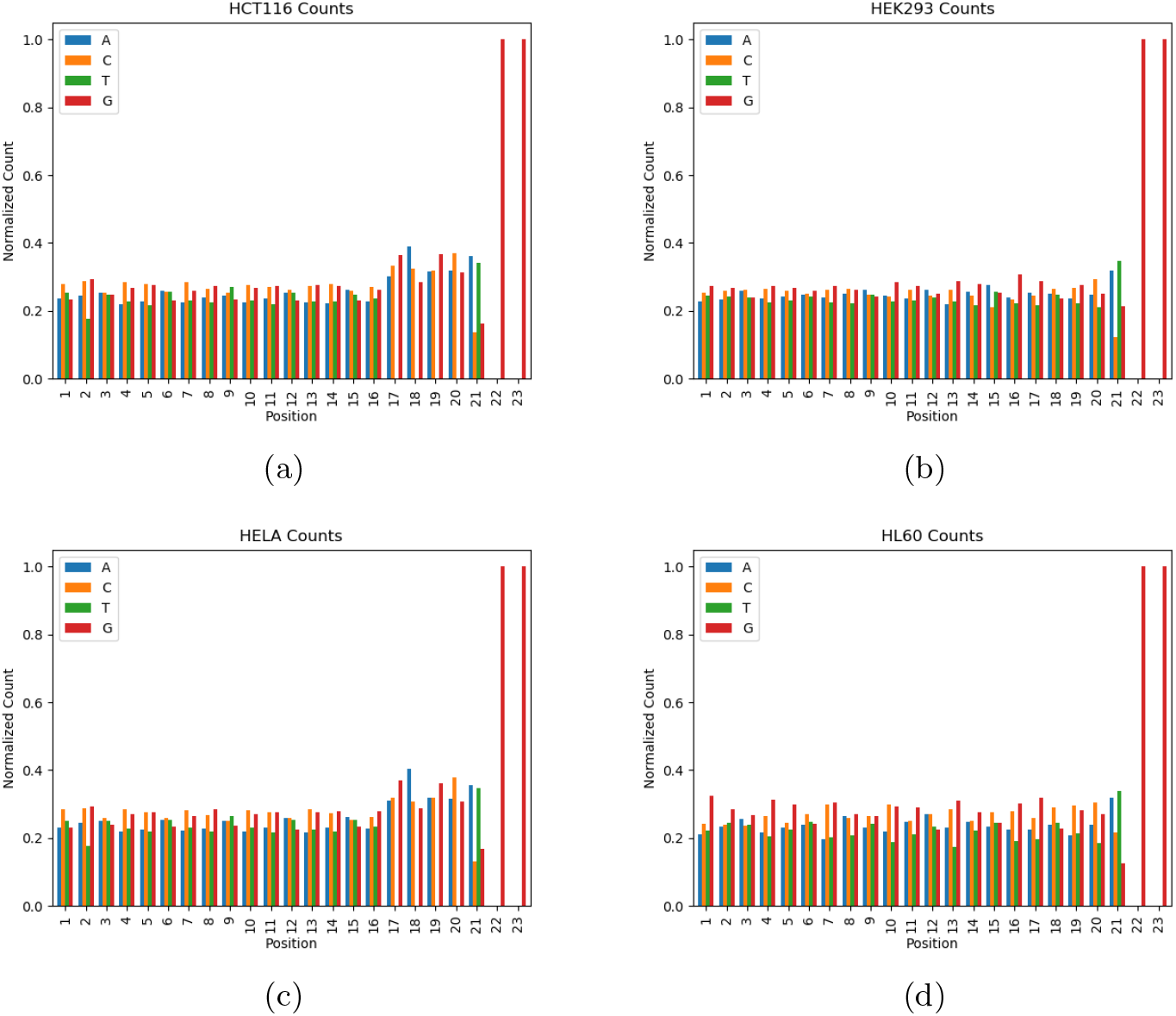
Position specific count of nucleotides for cell a) HCT116 b) HEK293 c) HeLa d) HL60. We can see from these figures that there is no Thymine at position 17, 18 and 19 for sgRNAs of cell type HCT116 and HeLa. But Thymine appears at these positions for sgRNAs of cell HEK293 and HL60

### 4.2 Feature Extraction

sgRNA sequence consists of four types of nucleotides: Adenine (A), Cytosine (C), Thymine (T) and Guanine (G). We extracted three types of features related to the composition of nucleotides in the sgRNAs. Two types of features were already used in CRISPRpred [10]. We added a new type of feature called n-gapped dipeptide which was used in [34]. Nevertheless, the features are described below for the sake of completeness.

- **Position Independent Features (PIF):** These features represent the number of occurrences of a given n adjacent nucleotides in the entire sequence. We extracted PIF features for *n* = 1, 2, 3,4. For example, the feature named AC indicates how many times the sequence AC appears in a given sgRNA sequence. For the sgRNA

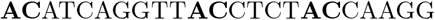

the number of times AC appears is 3. The number of PIF when *n* = 1 is 4, namely A, C, T, G. In the same way, number of PIF when *n* = 2 is 4^2^, i.e. AA, AC, …, GG. The number of PIF when *n* = 3 and *n* = 4 is 4^3^ and 4^4^ respectively. Thus, total number of PIF is

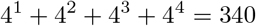
- **Position Specific Features (PSF):** This type of features are all binary features indicating whether a nucleotide or n adjacent nucleotides appear at a certain position in a given sgRNA. Again, we varied n from 1 to 4 for generating PSF. Feature named AA_2_ indicates whether the sequence AA appears at position 2 or not for a sgRNA. For the sgRNA

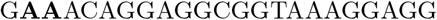

the value of AA_2_ is 1. When *n* = 1, there are 23 possible positions in which 4 possible nucleotides can appear. For *n* = 2, there are 22 possible positions in which 4^2^ possible 2 adjacent nucleotides can appear. In the same way, for *n* = 3, the number of possible position for 4^3^ possible 3 adjacent nucleotides to appear is 21. 4^4^ possible 4 adjacent nucleotides can appear at 20 possible positions. So, number of PSF, considering 1 ≤ *n* ≤ 4 is

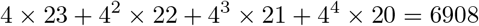
- ***n*-Gapped Dipeptides (nGD):** This type of features were used in [34]. In this type, we counted the number of times 2 given nucleotides appear at a certain distance in a sgRNA. The feature named GAP:AG_2_ specifies the number of times A and G occur at a distance of 2 nucleotides with order of nucleotides preserved. In other words, the value of GAP:AG_2_ may not be equal to the value of GAP:GA_2_. For the sgRNA,

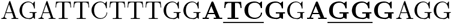

the value of GAP:AG_2_ is 2. 2 specific nucleotides can appear at a distance of 1, 2, 3,…, 21. Possible combination of 2 nucleotides is 4^2^. The total number of nGD is

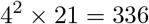

### 4.3 Feature Selection

We extracted a total of 7584 features. A large number of features have a chance of overfitting the model. Also, it increased the training time. To reduce the dimension of feature space, first we ranked the features using extremely randomized trees (ExtraTree) [35] using gini score. The number of estimator while training the (ExtraTree) was 500. Then we selected the features with gini scores greater than or equal to the mean gini score of all the features.

### 4.4 Training

#### 4.4.1 Standard Scaling

First we scaled all of our features using standard scaling. Standard scaling is a data pre-processing technique. It transforms data so that the mean of all data is 0 and standard deviation is 1. Among the extracted features there are binary features like if a nucleotide is present or absent at a specific position. This type of features can only be equal to 0 or 1. There are also some features like how many times a nucleotide occurs in the entire sequence. These features’ values have a different domain than binary features. So the feature values have varying ranges. There are several machine learning models that assume that the features have been scaled to the same value range. If there is a feature with a higher variance compared to other features then it will dominate the objective function and model will not be able to learn from other features.

#### 4.4.2 Support Vector Machine (SVM)

We experimented with several supervised machine learning algorithms to train our model. Finally non linear support vector machine (SVM) was used to train our final model. We used radial basis function (RBF) as the kernel for SVM. We also tuned the hyperparameters *C* and *γ* which has already been described. Finally, we trained our model for *C* =10 and *γ* = 0.001.

#### 4.4.3 Experimental Environment

We have conducted experiments using python language (version 3.6). We mainly used the scikit-learn package (version 0.20.3) [36] of python for all machine learning related programming. Pandas (0.24.2) and numpy (1.16.2) library was used for data management.

The experiments were carried out on Kaggle Kernel and in a server machine. Kaggle is a cloud computational environment. It had 4 CPU cores with 17 Gigabytes of RAM. In a single session it provides 9 hours of execution time with 5 Gigabytes of auto-saved disk space and 16 Gigabytes of temporary disk space. The server machine was equipped with Intel Xeon CPU E5-4617 @ 2.90GHz × 6, Ubuntu 13.04 64-bit OS, 15 MB L3 cache and 64 GB RAM.

## 5 Conclusion

CRISPRpred(SEQ) has performed exceptionally well and has almost beaten the state-of-the-art deep neural networking pipeline (i.e., DeepCRISPR) leveraging only traditional machine learning techniques and focusing only on primary sequence based features. In particular, CRISPRpred(SEQ) has improved upon the results of DeepCRISPR by 2.174%, 6.905% and 8.119% for the cells HCT116, HeLa and HL60 respectively. However, its performance for HEK293 cell is not up to the mark.

A future research direction from computational point of view is as follows. A prediction model can be built by combining the designed features and features extracted using deep learning methods. This may improve the performance of the prediction model.

All the cells of DeepCRISPR dataset except HEK293 are related to some form of cancer cell. This might be a reason behind bad performance in HEK293 cell. This needs further investigation perhaps from a biological angle.

